# Resurgence of a perinatal attraction for animate objects via thyroid hormone T_3_

**DOI:** 10.1101/2020.11.16.384289

**Authors:** Elena Lorenzi, Bastien S. Lemaire, Elisabetta Versace, Toshiya Matsushima, Giorgio Vallortigara

**Affiliations:** Center for Mind/Brain Sciences, University of Trento, Piazza Manifattura 1, 38068 Rovereto (TN), Italy; School of Biological and Chemical Sciences, Department of Biological and Experimental Psychology, Queen Mary University of London, London E1 4NS, United Kingdom; Department of Biology, Faculty of Science, Hokkaido University, N10-W8, Kita-Ku, Sapporo, Hokkaido 060-0810, Japan

## Abstract

For inexperienced brains, some stimuli are more attractive than others. Human neonates and newly-hatched chicks preferentially orient towards face-like stimuli, biological motion, and objects changing speed. In chicks, this enhances exposure to social partners, and subsequent attachment trough filial imprinting. Early preferences are not steady. The preference for stimuli changing speed fades away after three days in chicks. To understand the physiological mechanisms underlying these transient responses, we tested whether the early preferences for objects changing speed can be promoted by thyroid hormone 3,5,3’-triiodothyronine (T_3_). This hormone determines the start of imprinting’s sensitive period. We found that the preference for objects changing speed can be re-established in female chicks treated with T_3_. Moreover, day-one chicks treated with an inhibitor of endogenous T_3_ did not show any preference. These results suggest that the time windows of early predispositions and of high plasticity are controlled by the same molecular mechanisms.

## Introduction

Early experience plays a crucial role in shaping neural and behavioural development. However, the effects of early experience are stronger during certain periods of development (Hensch, 2005). An example is provided by filial imprinting (Bateson, 1979; Hess, 1959; Horn, 2004; McCabe, 2013; Spalding, 1873; Vallortigara & Versace, 2018). Shortly after birth or hatching, the young of some animals, usually of precocial species, learn to recognize their social partners (*e.g*., the mother and siblings) by simply being exposed to them. In the young domestic fowl (*Gallus gallus*), for instance, imprinting usually occurs within 24–48 h from hatching. Lorenz (1935) used the term “critical period” to refer to the fact that rather than being available throughout the lifespan, filial imprinting is shown only during a limited period of life. This time window can be extended for few more days, when a proper stimulus is not immediately available (Bolhuis, 1991). Being a flexible period, that has the characteristics of a self-terminating process (Bolhuis, 1991), imprinting is now better known as a “sensitive period”. Sensitive periods are windows of plasticity during brain development (Dehorter & Del Pino, 2020; Hensch, 2005).

The thyroid hormone 3,5,3’-triiodothyronine (T_3_) is implicated in the timing of the sensitive period. In domestic chicks, imprinting causes a rapid inflow of T_3_ in the brain (particularly the dorsal pallial regions including IMM, intermediate medial mesopallium) through converting enzyme (type 2 iodothyronine deiodinase, Dio_2_) localized in vascular endothelial cells in the brain (Yamaguchi et al., 2012) via Wnt-signaling pathway (Yamaguchi et al., 2018). Endogenous T_3_ level in the brain peaks around the peri-hatch period, and decays within a few days if not boosted by imprinting. However, in non-imprinted chicks whose sensitive period is ended, administration of exogenous T_3_ allows imprinting to occur, re-opening the sensitive period for memory formation (Yamaguchi et al., 2012) and the associated preference to biological motion (Miura et al., 2018). Re-opening of sensitive periods by pharmacological agents has been discovered for phenomena other than imprinting (see also Batista et al., 2016, 2018), such as ocular dominance in mammals (Hensch & Quinlan, 2018) and absolute pitch in humans (Gervain et al., 2013).

Although imprinting can be obtained with either naturalistic stimuli (resembling a conspecific) or artificial objects, a large amount of evidence shows the existence of spontaneous unlearned preferences for animate features (Rosa-Salva et al., under rev.). These preferences act as a sort of guidance or canalization mechanism to direct the newborns’ attention, favouring exposure to stimuli that are more likely to be social partners (reviewed in Di Giorgio et al., 2017; Rosa-Salva et al., under rev.; Versace et al., 2018). Preferences for animacy cues, that set apart animate from non-animate objects have been described in newly-hatched chicks, comprising *e.g*. preferences for face-like stimuli (Rosa-Salva et al., 2010), biological motion stimuli (Miura et al., 2020; Miura & Matsushima, 2016; Vallortigara et al., 2005) and self-propelled objects that move with variable speed (Rosa-Salva et al., 2016; for review see: Di Giorgio et al., 2017; Lorenzi & Vallortigara, in press). The same animacy cues operate on human newborns and other species (reviews: Di Giorgio et al., 2017; Lorenzi & Vallortigara, in press; Rosa-Salva et al., under rev.).

These biological priors, whose main function seems to speed up learning by canalizing imprinting, also operate during certain time periods only, revealing transient windows of sensitivity during development. Visually naïve chicks show a spontaneous preference for the head (face-like) region of a stuffed hen during the first two days post-hatching, which then fades away on day three (Johnson et al., 1989). Similarly, the spontaneous preference for objects moving with visible speed changes (Rosa-Salva et al., 2016) shows a window of sensitivity in three genetically selected and isolated breeds of chicks for only the first day of life, then disappearing on day three (Versace et al., 2019). Similarly, the biological motion preference occurs only within a few days post-hatch (Miura & Matsushima, 2012).

Here we report that the thyroid hormone T_3_ can modulate the timing of these windows of early preferences for animacy cues. These preferences, differently than the sensitive period for imprinting, do not depend on a particular experience with stimuli but rather guide attention towards particular stimuli, working as spontaneous, unlearned priors.

We devised three different experimental conditions. The first was aimed to confirm that the same window of sensitivity for the spontaneous animacy preference conveyed by visible speed changes shown in genetically-selected strains (Versace et al., 2019) also exists in the strain of broiler chicks we were using. We confirmed that the preference is there on post-hatching day 1, but fades away on day 3. To check whether T_3_ influences the duration of this window of sensitivity, we performed other two experiments. Firstly, to show that inhibition of endogenous T_3_ action can abolish the animacy preference we injected iopanoic acid (IOP), a potent inhibitor of the converting enzyme Dio_2_ that strongly reduces the brain T_3_ level (Yamaguchi et al., 2012) on post-hatching day 1, and compared IOP-injected chicks with vehicle-injected peers. Secondly, to show that the animacy preference can be re-established by exogenous T_3_ administration we injected chicks with T_3_ on post-hatching day 3 and compared T_3_-injected chicks with vehicle-injected peers.

## Results

The results for the not-injected condition are shown in Figure 1. The permutation test revealed a significant main effect of testing Day (*F*_(1, 67)_ = 5.40, *p* < 0.05), but did not reveal any effect of Sex (*F*_(1, 67)_ = 1.17, *p* = 0.28) nor interaction (*F*_(1, 67)_ = 0.05, *p* = 0.83; see Table 1 for the number of subjects tested in each condition for each sex, and for results split by sex). Chicks tested on day 1 showed a significant preference for animacy *V*_(35)_ = 458, *p* < 0.05, *d* = 0.44), whereas chicks tested on day 3 did not show any preference (*V*_(36)_ = 310, *p* = 0.72, *d* = 0.08).

**Figure 1:**
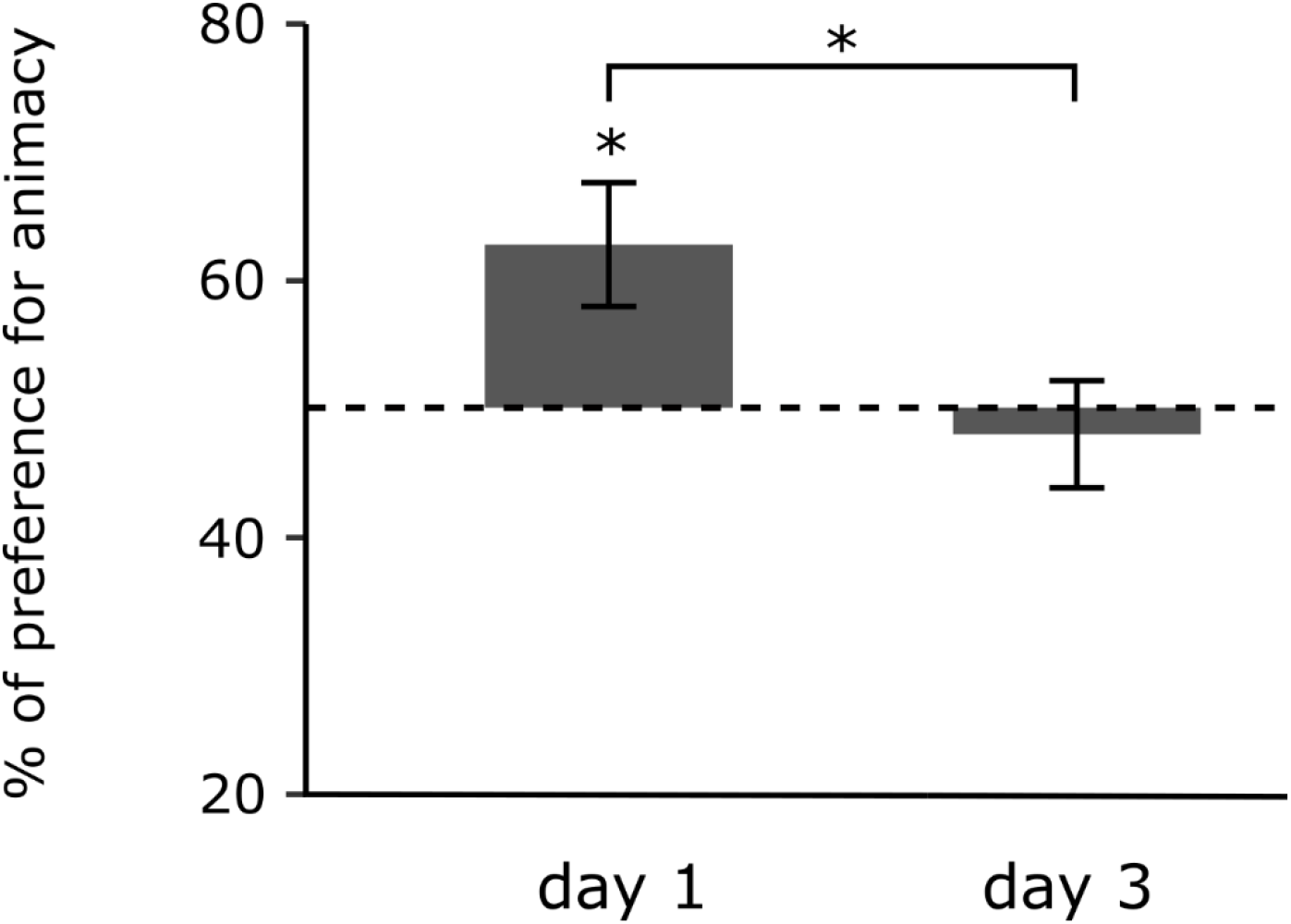
Preference for animacy for the not-injected chicks tested one or three days after hatching. Data are represented as mean ± SEM. Asterisks indicate significant differences from chance (black dotted line). Black asterisk between the two groups indicates a significant difference between testing Days. * indicates p<0.05.

**Table 1:** Summary of subjects tested and animacy preference in each condition. Number of subjects tested in each condition (N), subdivided by sex, mean preference for animacy, standard deviation (SD) and standard error (SE).

| Not-injected |  | DAY 1 |  |  |  | DAY 3 |  |  |  |
| --- | --- | --- | --- | --- | --- | --- | --- | --- | --- |
|  |  | N | Mean | SD | SEM | N | Mean | SD | SE |
|  | Females | 18 | 58.76 | 33.44 | 7.88 | 18 | 45.08 | 23.61 | 5.56 |
|  | Males | 17 | 67.20 | 23.45 | 5.69 | 18 | 50.71 | 27.45 | 6.47 |

| IOP-injected |  | IOP |  |  |  | Vehicle |  |  |  |
| --- | --- | --- | --- | --- | --- | --- | --- | --- | --- |
|  |  | N | Mean | SD | SEM | N | Mean | SD | SE |
| DAY 1 | Females | 16 | 48.36 | 29.55 | 7.39 | 14 | 69.40 | 19.29 | 5.15 |
|  | Males | 14 | 60.52 | 26.91 | 7.19 | 14 | 70.83 | 21.21 | 5.67 |

| T<sub>3</sub>-injected |  | T<sub>3</sub> |  |  |  | Vehicle |  |  |  |
| --- | --- | --- | --- | --- | --- | --- | --- | --- | --- |
|  |  | N | Mean | SD | SEM | N | Mean | SD | SE |
| DAY 3 | Females | 14 | 67.36 | 27.57 | 7.37 | 15 | 27.33 | 22.67 | 5.85 |
|  | Males | 15 | 37.65 | 23.31 | 6.02 | 17 | 62.94 | 27.62 | 6.70 |

The results for chicks treated on day 1 with the IOP (Dio_2_-inhibitor) are shown in Figure 2. The permutation test revealed a significant main effect of Treatment (*F*_(1,54)_ = 5.74, *p* < 0.05), but did not reveal any effect of Sex (*F*_(1, 54)_ = 1.10, *p* = 0.30) nor interaction (*F*_(1,67)_ = 0.66, *p* = 0.42; see Table 1). The vehicle-injected group showed a significant preference for animacy (*V*_(28)_ = 347, *p* < 0.001, *d* = 1.01), whereas the IOP-injected group did not show any preference (*V*_(30)_ = 268.5, *p* = 0.46, *d* = 0.14).

**Figure 2:**
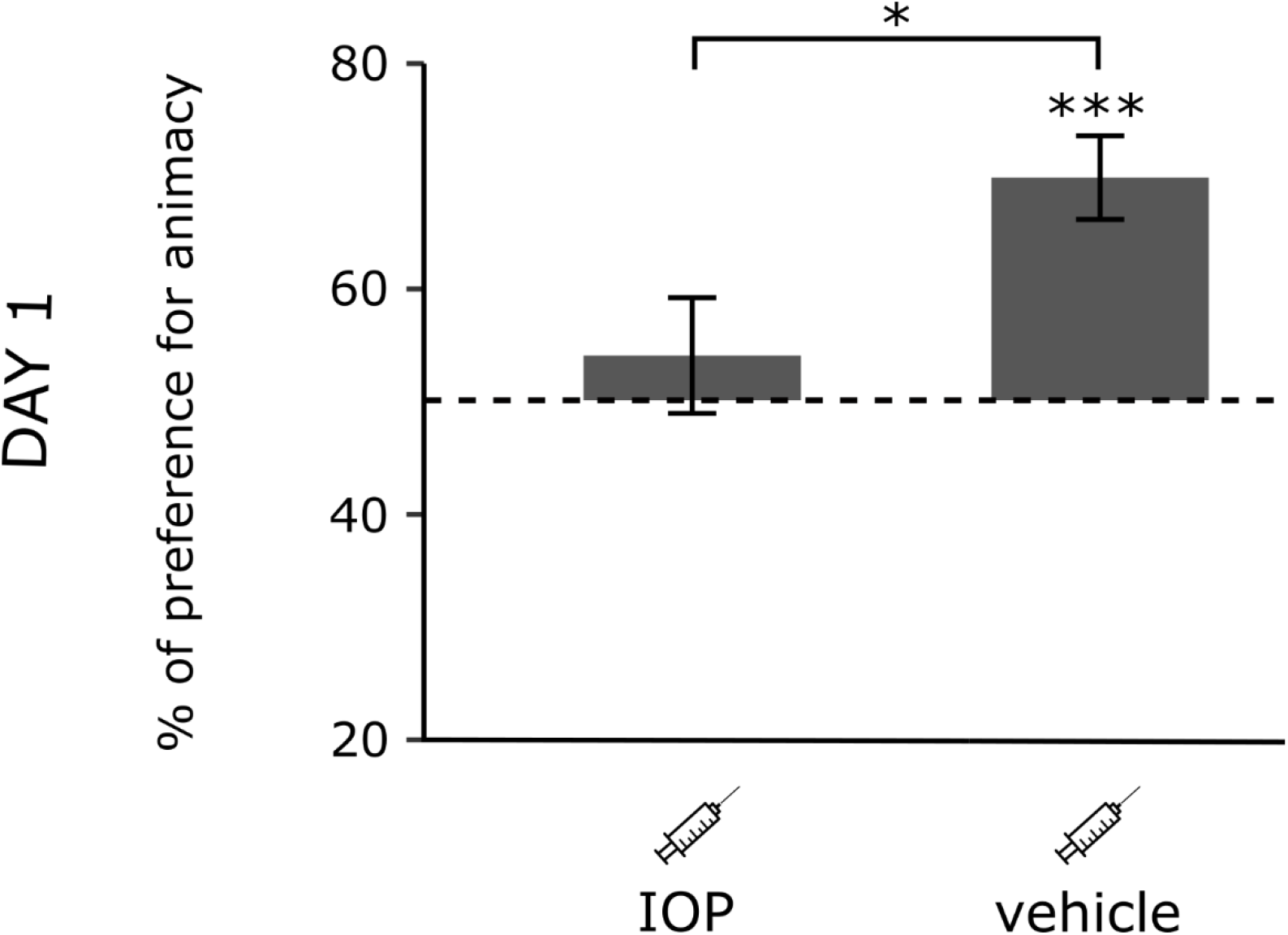
Preference for animacy for the chicks tested one day after hatching injected with IOP or with vehicle. Data are represented as mean ± SEM. Asterisks indicate significant differences from chance (black dotted line). Black asterisk between the two groups indicates a significant difference between IOP- and vehicle-injected chicks. *** indicates p<0.001, * indicates p<0.05.

The results for the chicks injected with T_3_ on day 3 are shown in Figure 3. The permutation test revealed a significant interaction between Treatment and Sex (*F*_(1,57)_ = 25.01, *p* < 0.001), but did not reveal any main effect of Treatment (*F*_(1, 57)_ = 1.27, *p* = 0.26) or Sex (*F*_(1, 57)_ = 0.20, *p* = 0.65). Females and males showed an opposite pattern of preference within each group, T_3_-injected (*W*_(29)_ = 166.5, *p* < 0.01, *d* = 1.17) and vehicle-injected (*W*_(32)_ = 39.5, *p* < 0.001, *d* = 1.40). T_3_- and vehicle-injected females showed a significant difference in their preferences (*W*_(28)_ = 183.5, *p* < 0.001, *d* = 1.59). T_3_-injected females showed a significant preference for animacy (*V*_(14)_ = 85, *p* < 0.05, *d* = 0.63), whereas vehicle-injected females showed a significant preference for the non-animacy stimulus (*V*_(15)_ = 11, *p* < 0.01, *d* = 1.0). T_3_- and vehicle-injected males also showed a significant difference in their preferences (*W*_(31)_ = 61, *p* < 0.05, *d* = 0.98). However, T_3_-injected males showed a marginally non-significant preference for the non-animacy stimulus (*V*_(15)_ = 27, *p* = 0.06, *d* = 0.53), whereas vehicle-injected males showed a marginally non-significant preference for animacy (*V*_(17)_ = 116, *p* = 0.06, *d* = 0.47).

**Figure 3:**
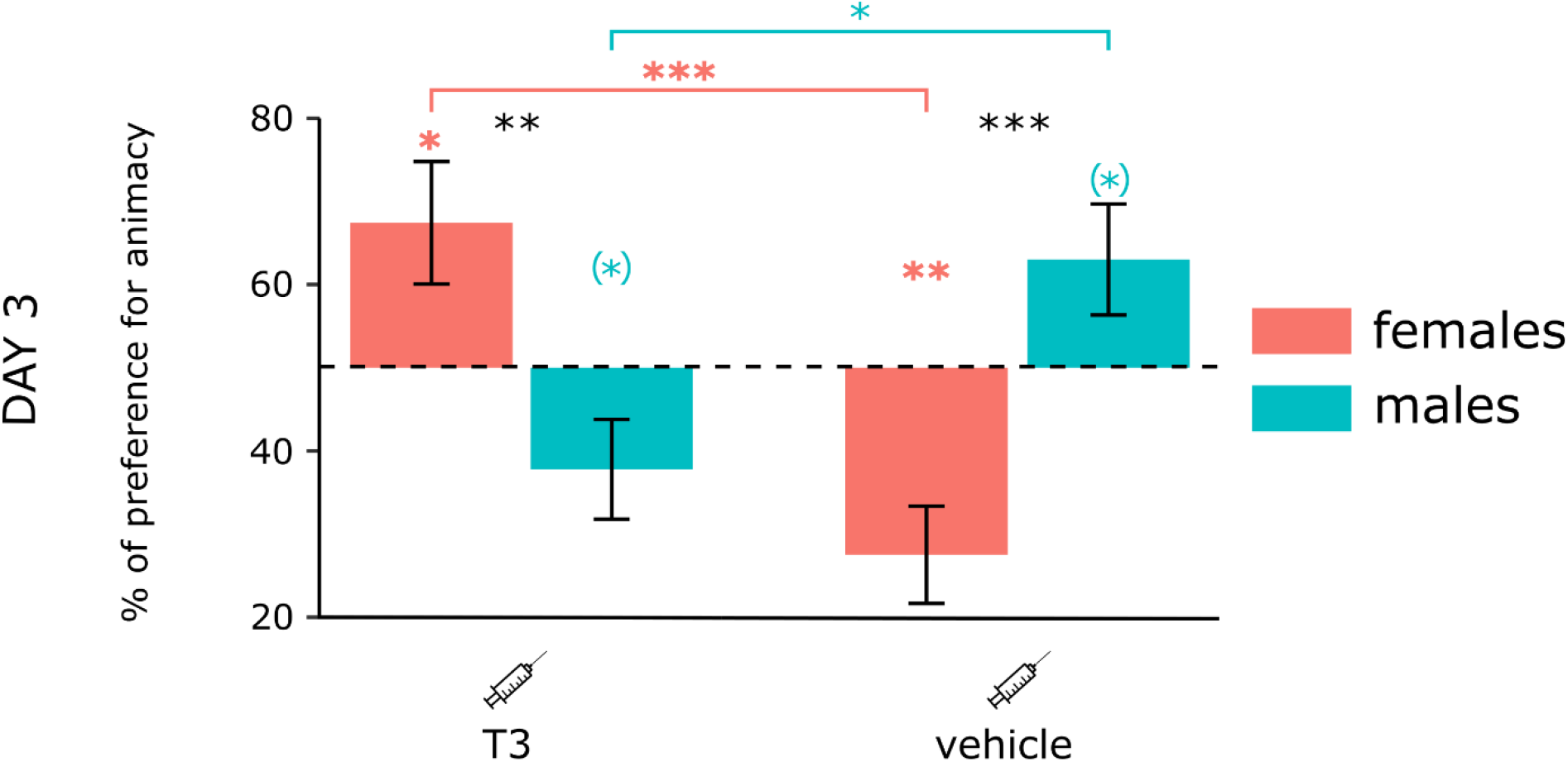
Preference for animacy for the chicks tested three days after hatching injected with T_3_ or vehicle. Data are represented as mean ± SEM. Black asterisks indicate a significant difference between sexes (females red and males blue) within each injection group. Red asterisks above a red horizontal line indicate a significant difference between females of each injection group. Blue asterisks above a blue horizontal line indicate a significant difference between males of each injection group. Red asterisks indicate a significant difference in females from chance. Blue asterisks show a difference from chance-level in males. *** indicates p<0.001, ** indicates p<0.01, * indicates p<0.05, (*) indicates p<0.07.

## Discussion

Untreated chicks showed a clear spontaneous preference for visible speed changes, a self-propelled cue to animacy (Rosa-Salva et al., 2016), on post-hatching day 1 that disappeared on day 3. Restricted sensitivity windows to animacy motion cues seem thus to exist in chicks similar to those for the head region of the mother hen (Johnson et al., 1989). Similar sensitivity windows have been described in humans as well (Buiatti et al., 2019; Johnson et al., 1991). During the first month of life infants preferentially look at schematic face-like configurations over identical stimuli in a non-face configuration, whereas the same is not observed at 3 and 5 months of age (Johnson et al., 1991). Similarly, the intensity of selective EEG responses to face-like patterns tends to decrease over time after the first hours of life (Buiatti et al., 2019).

The thyroid hormone 3,5,3’-triiodothyronine (T_3_) appears to play a crucial role in the sensitivity window for spontaneous motion animacy preference. Chicks injected on day 1 with an inhibitor of the converting enzyme (iopanoic acid, IOP) showed a complete lack of preference when compared with vehicle-injected chicks. Administration of T_3_ on day 3 seems to restore animacy preference, at least in females. Sex differences, which are well-known to be associated with critical period regulation by thyroid hormones (see (Batista & Hensch, 2019), seem to be complicated at this age also by the effect of injection in itself (even with vehicle only). T_3_-injected females preferred the animate stimulus, while vehicle-injected females preferred the non-animate stimulus. On the contrary, T_3_-injected males tended to prefer the non-animate stimulus, while vehicle-injected males tended to prefer the animate stimulus.

Precocious sex differences in social motivation and aggression are commonly observed in chicks (Andrew, 1966). Functionally, it has been argued that they may arise from different levels of social motivation (Cailotto et al., 1989; Vallortigara et al., 1990) and attitudes toward novelty (Vallortigara, 1992) in the two sexes. Females tend to engage more than males in social reinstatement and usually prefer to approach familiar individuals (Cailotto et al., 1989; Vallortigara, 1992; Vallortigara et al., 1990), whereas males tend to approach unfamiliar ones (Santolin et al., 2020; Vallortigara, 1992; Versace et al., 2017). In natural populations fowls exhibit territorial behaviour wherein single dominant cocks maintain and patrol a large territory within which a number of highly social-aggregated females live (McBride et al., 1969; McBride & Foenander, 1962). This sort of social organization may favour the prevalence of gregarious and affiliative behaviours in females and of aggressive and exploratory behaviours in males. Therefore, it could be that the injection of T_3_ affected both sexes similarly, but that they exhibited different behavioural responses coherent with their natural preferences that become apparent from day 3. Females would thus preferentially approach the template represented by the animacy stimulus, engaging more in social reinstatement responses, whereas males which are less socially motivated would have approached more the unfamiliar stimulus, engaging in exploratory and aggressive responses.

Note that vehicle-injected females on day 3 preferred the non-animacy stimulus, whereas males had a trend to prefer the animacy one. On day 3, chicks exhibit a peak of fear-related to hormonal levels (Schaller & Emlen, 1962). Males show a weaker avoidance response than females until post-hatching day 4. At this timepoint, sex differences observed in the vehicle group might arise from the different reactions to the stressful event provided by the needle. The fear caused by the injection in females might have evoked a fear response toward the most animate object, the animacy stimulus, making them approach the less animate one. In males, the same fear could have evoked an antipredator response enhancing social reinstatement towards the animacy stimulus.

Moving from the functional level to mechanisms, it is worth noticing that early during development the levels of different thyroid hormones (T_3_ and T_4_, thyroxine) in the brain show sex-dependent differences. The onset of the surge of T_4_ in male zebra finches (*Taeniopygia guttata)* precedes that of females, while the onset of T_3_ in females precedes that of males (Yamaguchi et al., 2017). Strengthening the hypothesis of an interaction between T_3_ and other sexually dimorphic hormones, response to animacy in visually naïve chicks involves brain regions rich in sex steroid hormone receptors (Lorenzi et al., 2017; Mayer et al., 2016; Mayer, Rosa-Salva, & Vallortigara, 2017; Mayer, Rosa-Salva, Morbioli, et al., 2017), part of the so-called *Social Behavior Network*, which appears to be highly conserved in vertebrates (Goodson & Kingsbury, 2013; Lorenzi et al., 2017; Newman, 1999; O’Connell & Hofmann, 2011).

Thyroid hormones are key regulators of vertebrates’ brain development (Van Herck et al., 2013). Among others, T_3_ directly influences the expression of genes by binding to its nuclear receptors, which act as transcription factors (Harvey & Williams, 2002). Therefore, abnormalities in the level of T_3_ during development may result in permanent impairments. In the present study, inhibiting T_3_ function gave rise to a lack of the spontaneous approach behaviour toward animate stimuli. Abnormalities in spontaneous preferences for animacy at birth have been linked to autistic spectrum disorder (Lorenzi et al., 2019; Sgadò et al., 2018). Interestingly, human neonates at high familial risk for autism exhibit anomalous looking patterns to animacy cues provided by schematic face-like and biological motion stimuli (Di Giorgio et al., 2016). Altogether, these results may point toward a T_3_ involvement in autistic spectrum disorders, paving the way for future investigations in humans.

In conclusion, we showed that providing exogenous T_3_ restores the predisposition to orient towards motion-animacy cue on day 3 post hatch and inhibiting the endogenous T_3_ conversion prevents the predisposition to appear on day 1 post-hatch. It is also notable that expression of Dio_2_ (the converting enzyme) is positively correlated with the imprinting-associated biological motion preference (Takemura et al., 2018). The similarity of effects of the thyroid hormone on imprinting and early predispositions is unlikely to be coincidental, for it would appear of little usage the re-opening of neural plasticity to the effects of environmental stimuli without restoring at the same time the biological priors to maximize the likelihood of getting proper environmental stimuli. The two processes should be functionally and temporally linked, and probably for this reason they are implemented in a similar molecular ground.

### Limitations of the Study

Here we considered one specific predisposition towards animate stimuli, relative to speed changes of motion stimuli. In order to show for the generality of the T_3_ effects on inborn animacy predispositions, others visual preferences such as those for face-like stimuli and biological motion need to be investigated.

### Resource Availability

▪ Further information and requests for resources and reagents should be directed to and will fulfilled by the Lead Contact, Dr. Elena Lorenzi
▪ The published article includes all datasets generated in this study

## Methods

### Animals

All experiments were carried out in compliance with the applicable European Union and Italian laws, and guidelines for animals’ care and use. All the experimental procedures were approved by the Ethical Committee of the University of Trento OPBA and by the Italian Health Ministry (permit number 1139/2015).

Fertilised eggs of the Aviagen Ross 308 strain were provided by a commercial hatchery (Azienda Agricola Crescenti, BS, Italy). Upon arrival, eggs were incubated under controlled temperature (37.7°C until post-hatching day 1, 33°C until post-hatching day 3) and humidity (60%) within incubators (FIEM MG140/200 Rural LCD EVO). Incubators were kept in darkness, preventing the chicks from any visual experience during incubation and hatching prior testing. We employed a total of 190 domestic chicks (*Gallus gallus*; 95 females; the exact number of chicks in each condition and sex is shown in *Table 1*).

### Apparatus and Stimuli

We used the same apparatus and stimuli as in previous studies investigating motion-animacy preference in newly-hatched chicks (Lorenzi et al., 2017, 2019; Rosa-Salva et al., 2016, 2018). A detailed description of the apparatus, stimuli and procedure used can be found in Rosa-Salva et al., 2016 (Exp. 2). Briefly, the apparatus consisted of a white corridor (85 x 30x 30 cm, see *Figure 4*) with two opposite high-frequency monitor screens (ASUS MG248Q, 24’’, 120 Hz). Three areas subdivided the corridor: a central one (starting area: 45 cm long) and two lateral ones (choice areas: 20 cm long each). A small step (1.5 cm high) on each side delimited the boundaries between starting and choice areas. A video-camera, centrally located above the apparatus, recorded animals’ behaviour. The two stimuli were displayed on the screens and represented a red circle (diameter 3 cm) moving horizontally. One stimulus was moving at a constant speed (≈4.64 cm/s on our monitors) while the other one was visibly changing speed (the slower speed being ≈3.37 cm/s and the faster one being ≈19.64 cm/s), a reliable cue to animacy (Rosa-Salva et al., 2016). We counterbalanced the position of the stimuli on the screens between subjects.

**Figure 4:**
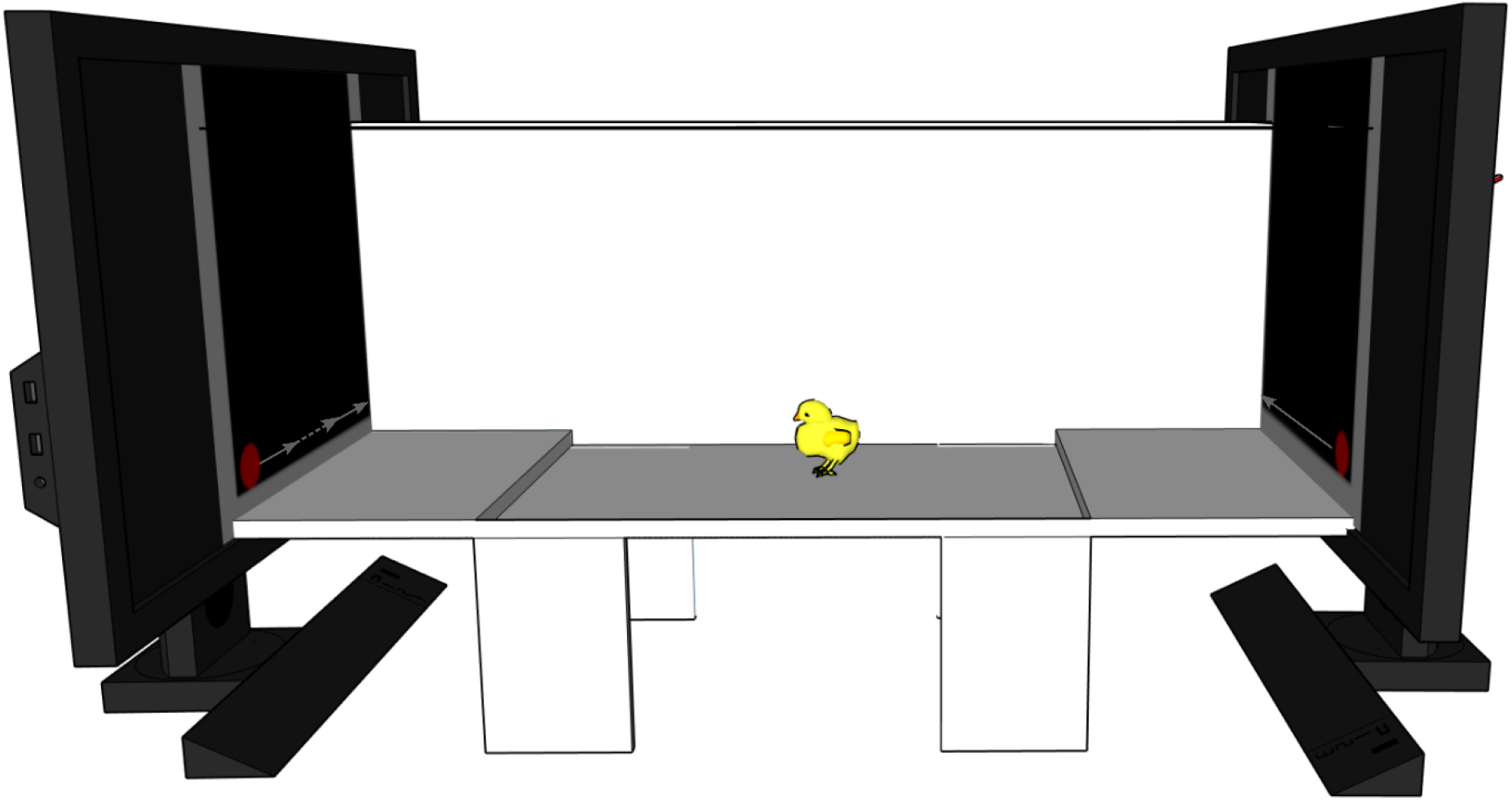
Schematic representation of the experimental apparatus. A white corridor with two video-screens at the two opposite ends playing the stimuli. On the left monitor is depicted the stimulus changing speed (dotted line represents faster speed), on the right monitor is represented the constant moving stimulus. The chick is represented in the starting area at the centre. Two little steps divide it from the two choice areas. For representative purposes one of the two longitudinal walls is represented translucent.

### Intramuscular injections

Iopanoic acid (IOP 10mM, TCI I0300, Tokyo Chemical Industry co. Ltd., Tokyo, Japan) was dissolved in 0.05M NaOH solution at 1mM and rebuffered to pH=8.5 by 6M HCl. 3,3’,5-Triiodo-L-thyronine (T3, 100 μM, Sigma Aldrich, T-2877) was dissolved in 0.002M NaOH and 0.9% NaCl. Vehicle solutions were also prepared to control for any effect of the injections. Respectively, the vehicle for IOP was a 0.05M NaOH solution buffered to pH=8.5 by 6M HCl, while the vehicle for T3 was a 0.9% NaCl and 0.002M NaOH solution (vehicle-injected groups). One hour before testing, each subject was carefully taken from the incubator in complete darkness. A black hood on the head prevented any source of visual stimulation during the intramuscular injection to thigh. Control chicks underwent the same procedure without receiving any injection (not-injected groups). Immediately after, each chick was placed back to the dark incubator. To distinguish single individuals in the darkness while keeping the same auditory environment experienced before injection, we placed the chicks in individual compartments within the same incubator.

### Testing

One hour after injection (according to the different groups assigned), each chick was individually tested for the spontaneous preference for animacy for 10 minutes. After placing each subject in the starting area, the time spent in each sector of the corridor was recorded. In order to measure the animal preference for motion-animacy, we considered the ratio [%] of time spent near the animacy stimulus over the total choice time using the formula:

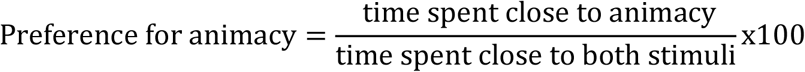

The preference score could range from 0% (full preference for the non-animacy stimulus) to 100% (full preference for animacy), while 50% represented the absence of preference. Permanence in the starting area for the total duration of the test was considered as no choice and led to the exclusion from further analyses (this occurred in about 47% of the cases). Sexing of the subjects occurred at the end of the procedure of testing.

### Statistical Analysis

The statistical analyses were performed using RStudio v1.1 and the following packages: *goftest* (Faraway et al., 2019), *nlme* (Pinheiro et al., 2020), *tidyr* (Wickham et al., 2020), *plyr* (Wickham, 2011), *dplyr* (Wickham et al., 2020), *reshape* (Wickham, 2007), *lsr* (Navarro, 2015), *ggplot2* (Wickham, 2016), *lmPerm* (Wheeler & Torchiano, 2016).

The number of subjects required in each group was a priori determined with a power analysis (Champely et al., 2020) with an effect size of 0.75, and an alpha of 0.05. Results showed that 16 individuals were required per group and per sex to achieve a power of 0.80.

To detect the presence of outliers, we used a Multivariate Model Approach using Cook’s distance. Subjects having a Cook’s distance 4 times greater than the group mean were considered as outliers and discarded from further analyses (Kannan & Manoj, 2015). We identified 6 outliers from different groups. To assess the normality of data distribution, we looked at the distribution of residuals (Q-Q plot).

As parametric assumptions were not met, we used non-parametric tests. In not-injected condition, to determine whether the testing Day (2 levels: day 1 and day 3) and Sex (2 levels: Female, Male) affected animacy preference, we performed a permutation test using F-test probabilities. A similar test was conducted for the IOP- and T_3_-injected conditions, to determine whether the treatments (2 levels: IOP/T_3_ and respective vehicles) and Sex (2 levels: Female, Male) affected animacy preference.

To determine whether the preference was statistically different between groups within each condition (not-, IOP-, T_3_-injected), we conducted two-sample Wilcoxon tests. To examine whether each group had a significant preference for either stimulus, we conducted one-sample Wilcoxon tests against chance level (50%). We calculated Cohen’s *d* (*d*) for each Wilcoxon test performed.

## Acknowledgments

This work was supported by a grant from the European Research Council under the European Union’s Seventh Framework Programme (FP7/2007-2013)/ Advanced Grant ERC PREMESOR G.A. [n 295517] to Giorgio Vallortigara. Authors are particularly thankful to Dr. Orsola Rosa-Salva for fruitful discussion.

## Author Contributions

Conceptualization, E.L., B.S.L., T.M., G.V. and E.V.; Methodology, E.L., B.S.L., T.M. and G.V.; Software, B.S.L.; Formal Analysis, E.L. and B.S.L.; Investigation, E.L. and B.S.L.; Resources, G.V. and T.M.; Writing-Original Draft, E.L. and B.S.L.; Writing-Review & Editing, G.V., T.M. and E.V.; Visualization, E.L. and B.S.L.; Funding Acquisition, G.V.

## References

Andrew, R. J. (1966). Precocious adult behaviour in the young chick. Animal Behaviour, 14(4), 485–500. https://doi.org/10.1016/s0003-3472(66)80050-7

Bateson, P. (1979). How do sensitive periods arise and what are they for? Animal Behaviour, 27, 470–486. https://doi.org/10.1016/0003-3472(79)90184-2

Batista, G., & Hensch, T. K. (2019). Critical Period Regulation by Thyroid Hormones: Potential Mechanisms and Sex-Specific Aspects. Frontiers in Molecular Neuroscience, 12. https://doi.org/10.3389/fnmol.2019.00077

Batista, G., Johnson, J. L., Dominguez, E., Costa-Mattioli, M., & Pena, J. L. (2016). Translational control of auditory imprinting and structural plasticity by eIF2α. ELife, 5, e17197. https://doi.org/10.7554/eLife.17197

Batista, G., Johnson, J. L., Dominguez, E., Costa-Mattioli, M., & Pena, J. L. (2018). Regulation of filial imprinting and structural plasticity by mTORC1 in newborn chickens. Scientific Reports, 8(1), 8044. https://doi.org/10.1038/s41598-018-26479-1

Bolhuis, J. J. (1991). Mechanisms of Avian Imprinting: A Review. Biological Reviews, 66(4), 303–345. https://doi.org/10.1111/j.1469-185X.1991.tb01145.x

Buiatti, M., Giorgio, E. D., Piazza, M., Polloni, C., Menna, G., Taddei, F., Baldo, E., & Vallortigara, G. (2019). Cortical route for facelike pattern processing in human newborns. Proceedings of the National Academy of Sciences, 116(10), 4625–4630. https://doi.org/10.1073/pnas.1812419116

Cailotto, M., Vallortigara, G., & Zanforlin, M. (1989). Sex differences in the response to social stimuli in young chicks. Ethology Ecology & Evolution, 1(4), 323–327. https://doi.org/10.1080/08927014.1989.9525502

Champely, S., Ekstrom, C., Dalgaard, P., Gill, J., Weibezahl, S., Anandkumar, A., & De Rosario, M. (2020). Package ‘pwr’.

Dehorter, N., & Del Pino, I. (2020). Shifting Developmental Trajectories During Critical Periods of Brain Formation. Frontiers in Cellular Neuroscience, 14. https://doi.org/10.3389/fncel.2020.00283

Di Giorgio, E., Frasnelli, E., Rosa-Salva, O., Maria Luisa, S., Puopolo, M., Tosoni, D., Simion, F., & Vallortigara, G. (2016). Difference in Visual Social Predispositions Between Newborns at Low-and High-risk for Autism. Scientific Reports, 6. https://doi.org/10.1038/srep26395

Di Giorgio, E., Loveland, J. L., Mayer, U., Rosa-Salva, O., Versace, E., & Vallortigara, G. (2017). Filial responses as predisposed and learned preferences: Early attachment in chicks and babies. Behavioural Brain Research Sree Test Content 1. https://doi.org/10.1016/j.bbr.2016.09.018

Faraway, J., Marsaglia, G., Marsaglia, J., & Baddeley, A. (2019). goftest: Classical Goodness-of-Fit Tests for Univariate Distributions (Version 1.2-2) [Computer software]. https://CRAN.R-project.org/package=goftest

Gervain, J., Vines, B. W., Chen, L. M., Seo, R. J., Hensch, T. K., Werker, J. F., & Young, A. H. (2013). Valproate reopens critical-period learning of absolute pitch. Frontiers in Systems Neuroscience, 7. https://doi.org/10.3389/fnsys.2013.00102

Goodson, J. L., & Kingsbury, M. A. (2013). What’s in a name? Considerations of homologies and nomenclature for vertebrate social behavior networks. Hormones and Behavior, 64(1), 103–112. https://doi.org/10.1016/j.yhbeh.2013.05.006

Harvey, C. B., & Williams, G. R. (2002). Mechanism of Thyroid Hormone Action. Thyroid, 12(6), 441–446. https://doi.org/10.1089/105072502760143791

Hensch, T. K. (2005). Critical period plasticity in local cortical circuits. Nature Reviews Neuroscience, 6(11), 877–888. https://doi.org/10.1038/nrn1787

Hensch, T. K., & Quinlan, E. M. (2018). Critical periods in amblyopia. Visual Neuroscience, 35, E014. https://doi.org/10.1017/S0952523817000219

Hess, E. H. (1959). Imprinting. Science, 130(3368), 133–141. JSTOR.

Horn, G. (2004). Pathways of the past: the imprint of memory. Nature Reviews Neuroscience, 5(2), 108–120. https://doi.org/10.1038/nrn1324

Johnson, M. H., Davies, D. C., & Horn, G. (1989). A sensitive period for the development of a predisposition in dark-reared chicks. Animal Behaviour, 37, 1044–1046. https://doi.org/10.1016/0003-3472(89)90148-6

Johnson, M. H., Dziurawiec, S., Ellis, H., & Morton, J. (1991). Newborns’ preferential tracking of face-like stimuli and its subsequent decline. Cognition, 40(1), 1–19. https://doi.org/10.1016/0010-0277(91)90045-6

Kannan, K. S., & Manoj, K. (2015). Outlier detection in multivariate data. Applied Mathematical Sciences, 9, 2317–2324. https://doi.org/10.12988/ams.2015.53213

Lorenz, K. (1935). Der Kumpan in der Umwelt des Vogels. Journal für Ornithologie, 83(3), 289–413. https://doi.org/10.1007/BF01905572

Lorenzi, E., Mayer, U., Rosa-Salva, O., & Vallortigara, G. (2017). Dynamic features of animate motion activate septal and preoptic areas in visually naïve chicks (Gallus gallus). Neuroscience, 354, 54–68. https://doi.org/10.1016/j.neuroscience.2017.04.022

Lorenzi, E., Pross, A., Rosa-Salva, O., Versace, E., Sgadò, P., & Vallortigara, G. (2019). Embryonic Exposure to Valproic Acid Affects Social Predispositions for Dynamic Cues of Animate Motion in Newly-Hatched Chicks. Frontiers in Physiology, 10. https://doi.org/10.3389/fphys.2019.00501

Lorenzi, E., & Vallortigara, G. (in press). Evolutionary and Neural Bases of the Sense of Animacy. In A. B. Kaufman, J. Call, & J. C. Kaufman (Eds.), Cambridge Handbook of Animal Cognition. Cambridge University Press.

Mayer, U., Rosa-Salva, O., Lorenzi, E., & Vallortigara, G. (2016). Social predisposition dependent neuronal activity in the intermediate medial mesopallium of domestic chicks (Gallus gallus domesticus). Behavioural Brain Research, 310, 93–102. https://doi.org/10.1016/j.bbr.2016.05.019

Mayer, U., Rosa-Salva, O., Morbioli, F., & Vallortigara, G. (2017). The motion of a living conspecific activates septal and preoptic areas in naive domestic chicks (Gallus gallus). European Journal of Neuroscience, 45(3), 423–432. https://doi.org/10.1111/ejn.13484

Mayer, U., Rosa-Salva, O., & Vallortigara, G. (2017). First exposure to an alive conspecific activates septal and amygdaloid nuclei in visually-naïve domestic chicks (Gallus gallus). Behavioural Brain Research, 317, 71–81. https://doi.org/10.1016/j.bbr.2016.09.031

McBride, G., & Foenander, F. (1962). Territorial Behaviour in Flocks of Domestic Fowls. Nature, 194(4823), 102–102. https://doi.org/10.1038/194102a0

McBride, G., Parer, I. P., & Foenander, F. (1969). The Social Organization and Behaviour of the Feral Domestic Fowl. Animal Behaviour Monographs, 2, 125–181. https://doi.org/10.1016/S0066-1856(69)80003-8

McCabe, B. J. (2013). Imprinting. Wiley Interdisciplinary Reviews: Cognitive Science, 4(4), 375–390. https://doi.org/10.1002/wcs.1231

Miura, M., Aoki, N., Yamaguchi, S., Homma, K. J., & Matsushima, T. (2018). Thyroid Hormone Sensitizes the Imprinting-Associated Induction of Biological Motion Preference in Domestic Chicks. Frontiers in Physiology, 9. https://doi.org/10.3389/fphys.2018.01740

Miura, M., & Matsushima, T. (2012). Preference for biological motion in domestic chicks: sex-dependent effect of early visual experience. Animal Cognition, 15(5), 871–879. https://doi.org/10.1007/s10071-012-0514-x

Miura, M., & Matsushima, T. (2016). Biological motion facilitates filial imprinting. Animal Behaviour, 116, 171–180. https://doi.org/10.1016/j.anbehav.2016.03.025

Miura, M., Nishi, D., & Matsushima, T. (2020). Combined predisposed preferences for colour and biological motion make robust development of social attachment through imprinting. Animal Cognition, 23(1), 169–188. https://doi.org/10.1007/s10071-019-01327-5

Navarro, D. J. (2015). Learning Statistics with R: A tutorialfor psychology students and other beginners. (version 0.5). https://learningstatisticswithr.com

Newman, S. W. (1999). The Medial Extended Amygdala in Male Reproductive Behavior A Node in the Mammalian Social Behavior Network. Annals of the New York Academy of Sciences, 877(1), 242–257. https://doi.org/10.1111/j.1749-6632.1999.tb09271.x

O’Connell, L. A., & Hofmann, H. A. (2011). The Vertebrate mesolimbic reward system and social behavior network: A comparative synthesis. The Journal of Comparative Neurology, 519(18), 3599–3639. https://doi.org/10.1002/cne.22735

Pinheiro, J., Bates, D., to 2002), S. D. (up, Sarkar, D., & Heisterkamp, S. (2020). nlme: Linear and Nonlinear Mixed Effects Models (Version 3.1-148) [Computer software]. https://CRAN.R-project.org/package=nlme

Rosa-Salva, O., Fiser, J., Versace, E., Dolci, C., Chehaimi, S., Santolin, C., & Vallortigara, G. (2018). Spontaneous Learning of Visual Structures in Domestic Chicks. Animals: An Open Access Journal from MDPI, 8(8). https://doi.org/10.3390/ani8080135

Rosa-Salva, O., Grassi, M., Lorenzi, E., Regolin, L., & Vallortigara, G. (2016). Spontaneous preference for visual cues of animacy in naïve domestic chicks: The case of speed changes. Cognition, 157, 49–60. https://doi.org/10.1016/j.cognition.2016.08.014

Rosa-Salva, O., Mayer, U., Versace, E., Hebert, M., Lemaire, B. S., & Vallortigara, G. (under rev.). Sensitive periods for social development: Interactions between predisposed and learned mechanisms. Cognition.

Rosa-Salva, O., Regolin, L., & Vallortigara, G. (2010). Faces are special for newly hatched chicks: evidence for inborn domain-specific mechanisms underlying spontaneous preferences for face-like stimuli. Developmental Science, 13(4), 565–577. https://doi.org/10.1111/j.1467-7687.2009.00914.x

Santolin, C., Rosa-Salva, O., Lemaire, B. S., Regolin, L., & Vallortigara, G. (2020). Statistical learning in domestic chicks is modulated by strain and sex. Scientific Reports, 10(1), 15140. https://doi.org/10.1038/s41598-020-72090-8

Schaller, G. B., & Emlen, J. T. (1962). The ontogeny of avoidance behaviour in some precocial birds. Animal Behaviour, 10(3), 370–381. https://doi.org/10.1016/0003-3472(62)90060-X

Sgadò, P., Rosa-Salva, O., Versace, E., & Vallortigara, G. (2018). Embryonic Exposure to Valproic Acid Impairs Social Predispositions of Newly-Hatched Chicks. Scientific Reports, 8(1), 5919. https://doi.org/10.1038/s41598-018-24202-8

Spalding, D. (1873). Instinct, with original observations on young animals. Macmillan’s Mag, 27, 282–293.

Takemura, Y., Yamaguchi, S., Aoki, N., Miura, M., Homma, K. J., & Matsushima, T. (2018). Gene expression of Dio2 (thyroid hormone converting enzyme) in telencephalon is linked with predisposed biological motion preference in domestic chicks. Behavioural Brain Research, 349, 25–30. https://doi.org/10.1016/j.bbr.2018.04.039

Vallortigara, G. (1992). Affiliation and aggression as related to gender in domestic chicks (Gallus gallus). Journal of Comparative Psychology, 106(1), 53–57. https://doi.org/10.1037/0735-7036.106.1.53

Vallortigara, G., Cailotto, M., & Zanforlin, M. (1990). Sex differences in social reinstatement motivation of the domestic chick (Gallus gallus) revealed by runway tests with social and nonsocial reinforcement. Journal of Comparative Psychology (Washington, D.C.: 1983), 104(4), 361–367. https://doi.org/10.1037/0735-7036.104.4.361

Vallortigara, G., Regolin, L., & Marconato, F. (2005). Visually Inexperienced Chicks Exhibit Spontaneous Preference for Biological Motion Patterns. PLoS Biol, 3(7), e208. https://doi.org/10.1371/journal.pbio.0030208

Vallortigara, G., & Versace, E. (2018). Filial Imprinting. In J. Vonk & T. Shackelford (Eds.), Encyclopedia of Animal Cognition and Behavior (Springer).

Van Herck, S. L. J., Geysens, S., Delbaere, J., & Darras, V. M. (2013). Regulators of thyroid hormone availability and action in embryonic chicken brain development. General and Comparative Endocrinology, 190, 96–104. https://doi.org/10.1016/j.ygcen.2013.05.003

Versace, E., Martinho-Truswell, A., Kacelnik, A., & Vallortigara, G. (2018). Priors in Animal and Artificial Intelligence: Where Does Learning Begin? Trends in Cognitive Sciences. https://doi.org/10.1016/j.tics.2018.07.005

Versace, E., Ragusa, M., & Vallortigara, G. (2019). A transient time window for early predispositions in newborn chicks. Scientific Reports, 9(1), 18767. https://doi.org/10.1038/s41598-019-55255-y

Versace, E., Spierings, M. J., Caffini, M., Ten Cate, C., & Vallortigara, G. (2017). Spontaneous generalization of abstract multimodal patterns in young domestic chicks. Animal Cognition, 20(3), 521–529. https://doi.org/10.1007/s10071-017-1079-5

Wheeler, B., & Torchiano, M. (2016). lmPerm: Permutation Tests for Linear Models (Version 2.1.0) [Computer software]. https://CRAN.R-project.org/package=lmPerm

Wickham, H. (2007). Reshaping Data with the **reshape** Package. Journal of Statistical Software, 21(12). https://doi.org/10.18637/jss.v021.i12

Wickham, H. (2011). The Split-Apply-Combine Strategy for Data Analysis. Journal of Statistical Software, 40(1), 1–29. https://doi.org/10.18637/jss.v040.i01

Wickham, H. (2016). ggplot2: Elegant Graphics for Data Analysis. Springer.

Wickham, H., Henry, L., & RStudio. (2020). tidyr: Tidy Messy Data (Version 1.1.1) [Computer software]. https://CRAN.R-project.org/package=tidyr

Yamaguchi, S., Aoki, N., Kitajima, T., Iikubo, E., Katagiri, S., Matsushima, T., & Homma, K. J. (2012). Thyroid hormone determines the start of the sensitive period of imprinting and primes later learning. Nature Communications, 3, 1081. https://doi.org/10.1038/ncomms2088

Yamaguchi, S., Aoki, N., Matsushima, T., & Homma, K. J. (2018). Wnt-2b in the intermediate hyperpallium apicale of the telencephalon is critical for the thyroid hormone-mediated opening of the sensitive period for filial imprinting in domestic chicks (Gallus gallus domesticus). Hormones and Behavior, 102, 120–128. https://doi.org/10.1016/j.yhbeh.2018.05.011

Yamaguchi, S., Hayase, S., Aoki, N., Takehara, A., Ishigohoka, J., Matsushima, T., Wada, K., & Homma, K. J. (2017). Sex Differences in Brain Thyroid Hormone Levels during Early Post-Hatching Development in Zebra Finch (Taeniopygia guttata). PLOS ONE, 12(1), e0169643. https://doi.org/10.1371/journal.pone.0169643

